# Jasmonate Primes Plant Responses to Extracellular ATP

**DOI:** 10.1101/2024.11.07.622526

**Authors:** Jeremy B. Jewell, Ashleigh S. Carlton, Jordan P. Tolley, Laura E. Bartley, Kiwamu Tanaka

## Abstract

Extracellular ATP (eATP) signaling in *Arabidopsis thaliana* is mediated by the purinoceptor P2K1. Previous studies have clarified that the downstream transcriptional responses to eATP involve jasmonate (JA)-based signaling components such as the JA receptor (COI1) and JA-responsive bHLH transcription factors (MYCs). However, the specific contributions of JA signaling itself on eATP signaling are unexplored. Here, we report that JA primes plant responses to eATP through P2K1. Our findings show that JA treatment significantly upregulates *P2K1* transcription, corroborating our observation that JA facilitates eATP-induced cytosolic calcium elevation and transcriptional reprogramming in a JA signaling-dependent manner. Additionally, we find that salicylic acid pretreatment represses eATP-induced plant response. These results suggest that JA accumulation during biotic or abiotic stresses may potentiate eATP signaling, enabling plants to better cope with subsequent stress events.

**SIGNIFICANCE STATEMENT:** Plant hormone jasmonate (JA) enhances plant responses to extracellular ATP (eATP) in *Arabidopsis thaliana* through a mechanism dependent on the JA receptor COI1 and the eATP receptor P2K1. The reciprocal amplification of these signals provides a mechanistic explanation for how plants effectively respond to different stress events.

## INTRODUCTION

ATP is present in high concentrations within the cell (∼mM) but is typically found at much lower levels (∼nM) in the extracellular space, where it functions as a potent signaling molecule essential for cellular communication **(Watt et al., 1998; Weerasinghe et al., 2000)**. This purinergic signaling is well-documented across eukaryotic systems **(Burnstock and Verkhratsky 2009)**. In animals, extracellular ATP (eATP) has been extensively studied since the 1970s for its roles in neurotransmission, cellular physiology, and pathophysiology **(Burnstock 2012)**. More recently, purinergic signaling in plants has gained considerable attention, particularly following the identification of the first plant purinoceptor, the receptor kinase P2K1. This receptor was discovered through Arabidopsis mutants that do not respond to exogenous ATP treatment with the typical increase in cytosolic calcium levels ([Ca^2+^]_cyt_)—a characteristic of purinergic signaling **(Choi et al., 2014)**.

An emerging theme of these studies is the regulation of plant growth and defense mechanisms by P2K1 and eATP, especially under biotic and abiotic stresses **(Chen et al., 2017, Kumar er al. 2020; Jewell et al., 2022; Myers et al., 2022; Kim et al., 2023a; Kim et al., 2023b; Kim et al., 2023c; Sowders et al., 2023; Sowders et al.; 2024)**. Thus, eATP is recognized as a damage-associated molecular pattern (DAMP) in plants, triggering defense responses upon cellular injury (**Tanaka et al., 2014; Tanaka and Heil 2021**).

Transcriptional responses to eATP show remarkable similarity to those induced by wounding or treatment with the plant hormone jasmonic acid (JA) **(Choi et al., 2014; Jewell et al., 2019)**, an essential regulator of plant responses to various stresses **(Zhang et al., 2008; Browse 2009; Howe et al., 2018; Pascual et al., 2023)**. Similarly, eATP has also been implicated in systemic responses to leaf wounding **(Myers et al., 2022)**, a process largely mediated by JA signaling **(Reymond et al., 2000; Devoto et al., 2005)**. Notably, molecular components of the JA signaling pathway, including the receptor COI1 and bHLH transcription factors MYC2, MYC3, and MYC4, are required for eATP-induced transcriptional responses, indicating a significant interaction between eATP and JA signaling pathways **(Jewell et al., 2019)**.

While JA signaling components are essential for eATP responses, the influence of JA on eATP-mediated processes remains unexplored. Since eATP signaling depends on JA pathway components, understanding how JA affects eATP-induced responses is critical. Here, we investigate this relationship by examining how JA modulates eATP responses in *Arabidopsis thaliana*. Our findings indicate that JA primes the plant for eATP signaling by upregulating the *P2K1* receptor gene, suggesting a functional interplay in which JA signaling enhances plant sensitivity to eATP.

## RESULTS

### JAZ1 protein stability is not affected by eATP signaling

It has been demonstrated that eATP acts through JA signaling, as many eATP-responsive genes require the JA receptor COI1 for their expression **(Jewell et al., 2019)**. It has also been reported that eATP stimulus promotes the degradation of a jasmonate co-receptor protein (JAZ1) in a jasmonate-independent manner **(Tripathi et al., 2018)**. The mechanism of this surprising phenomenon has not been investigated in subsequent reports; therefore, we re-tested the effect of eATP on JAZ1 protein stability. Arabidopsis plants constitutively expressing the JAZ1-β-GLUCURONIDASE (JAZ1-GUS) fusion protein **(Thines et al., 2007)** were treated with 2 μM methyl jasmonate (MeJA) or 1 mM ATP for 30 minutes and JAZ1-GUS protein levels were quantified using a biochemical GUS activity assay **(Figure 1)**. Across three independent and not statistically different trials (*P=*0.13, one factor ANOVA F-test), MeJA treatment reduced JAZ1-GUS levels by approximately 48% (*P*<1e-7, Tukey HSD), while ATP treatment had no detectable effects (*P*=0.99, Tukey HSD). Similarly, ATP treatment did not affect JAZ1-GUS stability in the *aos* JA biosynthesis mutant (*P*=0.97, Tukey HSD), whereas MeJA still reduced GUS levels by about 49% (*P=*5.3e-3, Tukey HSD; **Supplemental Figure S1**). These findings suggest that while eATP signaling depends on COI1, it is independent of JAZ protein stability, indicating a distinct mechanism of JA-signaling-mediated eATP response, and suggesting that a model of eATP-stimulated JAZ1 degradation should be critically reevaluated.

**Figure 1.**
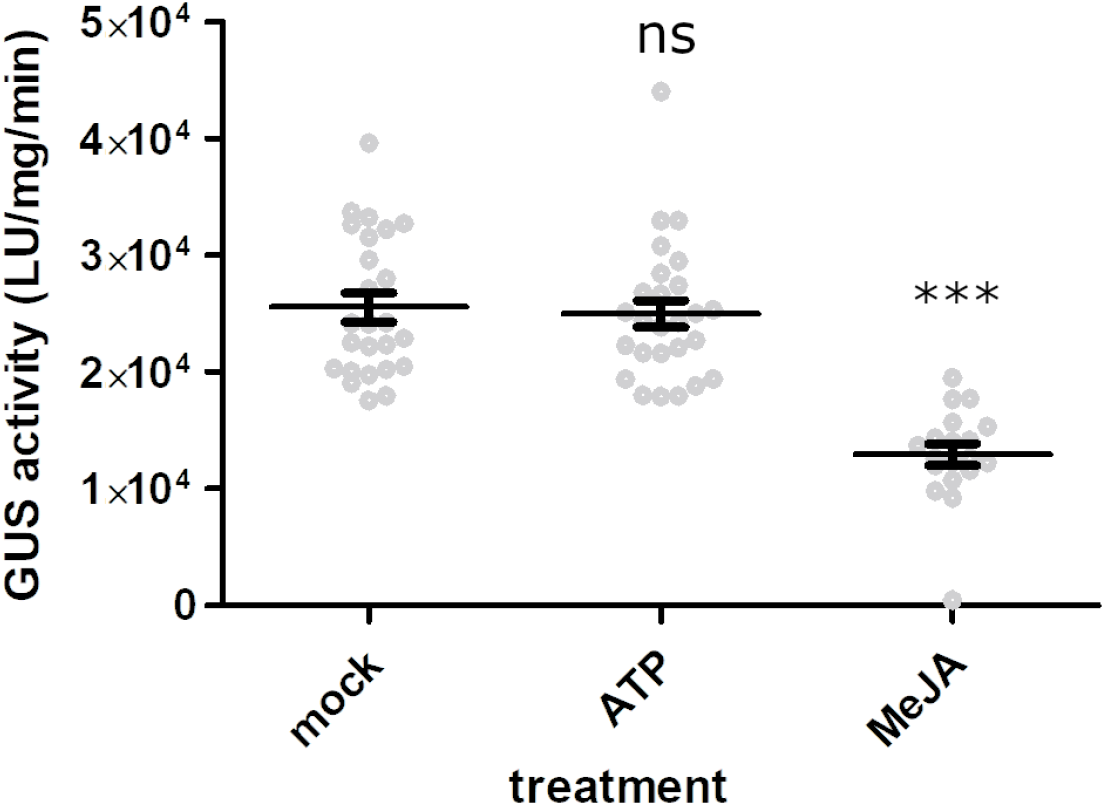
Exogenous ATP treatment does not reduce JAZ1 protein stability in *Arabidopsis* seedlings, while methyl jasmonate does. WT seedlings expressing JAZ1-GUS were treated with 2 μM MeJA or 1 mM ATP for 30 minutes and processed for quantitative GUS activity measurements to assess the JAZ1 protein stability, as described in Methods. Statistical comparisons to mock-treated seedlings are indicated by *** and ns for *P <* 0.001 and *P >* 0.99, respectively (Fisher’s LSD). Error bars indicate SEM, with *n* = 19-27 replicates per treatment across 3 trials.

### JA pre-treatment enhances eATP-induced cytosolic calcium response through a P2K1-dependent pathway

Given that eATP signaling integrates with JA pathways, we next explored whether JA itself modulates plant responses to eATP. To test this, we pre-treated transgenic Arabidopsis seedling expressing the calcium reporter apoaequorin protein (hereafter AEQ seedlings) overnight with 50 μM MeJA or a mock solution before adding 100 μM exogenous ATP to the same wells and recorded luminescence as an indication of cytosolic calcium levels ([Ca^2+^]_cyt_) as previously described (Choi et al., 2014). Without MeJA pre-treatment, ATP induced the typically transient [Ca^2+^]_cyt_ response **(Tanaka et al., 2010)**, peaking around ∼30-45 seconds and tapering thereafter **(Figure 2A)**. However, MeJA pre-treatment resulted in enhanced response to eATP, approximately 50% higher than in mock-treated samples over the course of 3 minutes and with a significantly earlier peak calcium level **(Figure 2, Supplemental Figure S2)**. We note that overnight pretreatment with 0.05% DMSO (the MeJA carrier) resulted in ∼18% reduced [Ca^2+^]_cyt_ response to ATP treatment **(Supplemental Figure S3)**, so the MeJA-primed eATP-induced calcium values are an underestimate. We also note that neither DMSO nor 50 μM MeJA treatment differed in changes to [Ca^2+^]_cyt_ relative to water **(Supplemental Figure S4)**. In contrast to WT seedlings, the *p2k1-3* mutant (lacking functional P2K1) did not show the typical [Ca^2+^]_cyt_ response to ATP **(Figure 2)**, regardless of MeJA pre-treatment, confirming that the JA-enhanced plant response to ATP is P2K1-dependent and the previously described second eATP receptor, P2K2 (Pham et al., 2020), is insufficient to allow for detectable changes to [Ca^2+^]_cyt_ in response to ATP. Interestingly, we were also able to detect MeJA-stimulated release of ATP into media surrounding leaf discs 1 hour and 16 hours after treatment, though not after 15 minutes **(Supplemental Figure S5)**. The mechanism and significance of this observation remains to be determined.

**Figure 2.**
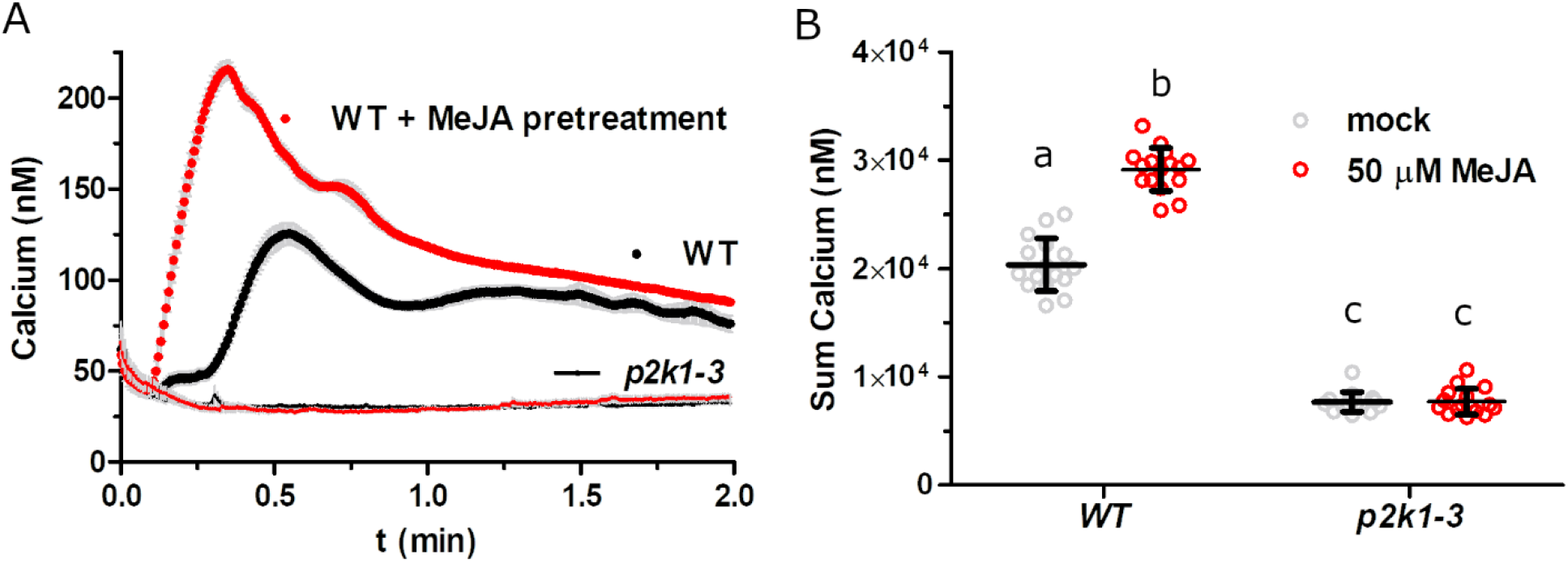
JA pre-treatment increases ATP-induced cytosolic calcium response. **(A)** Seven-day-old aequorin-expressing seedlings were incubated overnight in reconstitution buffer and treated with (red symbols) or without (black symbols) 50 μM MeJA, followed by treatment with 100 μM ATP. Aequorin luminescence was recorded for 2 minutes to assess the cytosolic calcium levels (mean ± SEM, *n* = 16 seedlings per treatment). **(B)** Summed calcium values from panel A. Different letters indicate statistically significant differences (*P* < 0.05, Tukey HSD). Mean, individual values, and SEM are shown, with *n* = 16 seedlings per treatment.

Next, we asked whether shorter MeJA pre-treatment durations could also enhance the ATP-induced [Ca^2+^]_cyt_ response. Replacing the medium with MeJA-containing solution at 18, 6, or 3 hours prior to ATP treatment revealed that even the shortest duration (3 hours) reliably increased the ATP-induced [Ca^2+^]_cyt_ response, with longer pre-treatment durations showing progressively larger effects **(Figure 3A,B**). Testing MeJA concentrations from 6.25 to 100 μM revealed a dose-dependent enhancement, where even the lowest concentration (6.25 μM) significantly increased ATP-induced [Ca^2+^]_cyt_ levels, with higher MeJA doses resulting in greater responses **(Figure 3C,D)**. One hour after wounding, Arabidopsis leaves contain JA at about 1-3 nanomoles per gram of fresh weight, or roughly micromolar concentration **(Kimberlin et al., 2022)**. Thus, these MeJA treatment dosages and durations are of the same magnitude of physiological JA levels after wounding, suggesting that JA priming may naturally augment eATP responses in wounded plants.

**Figure 3.**
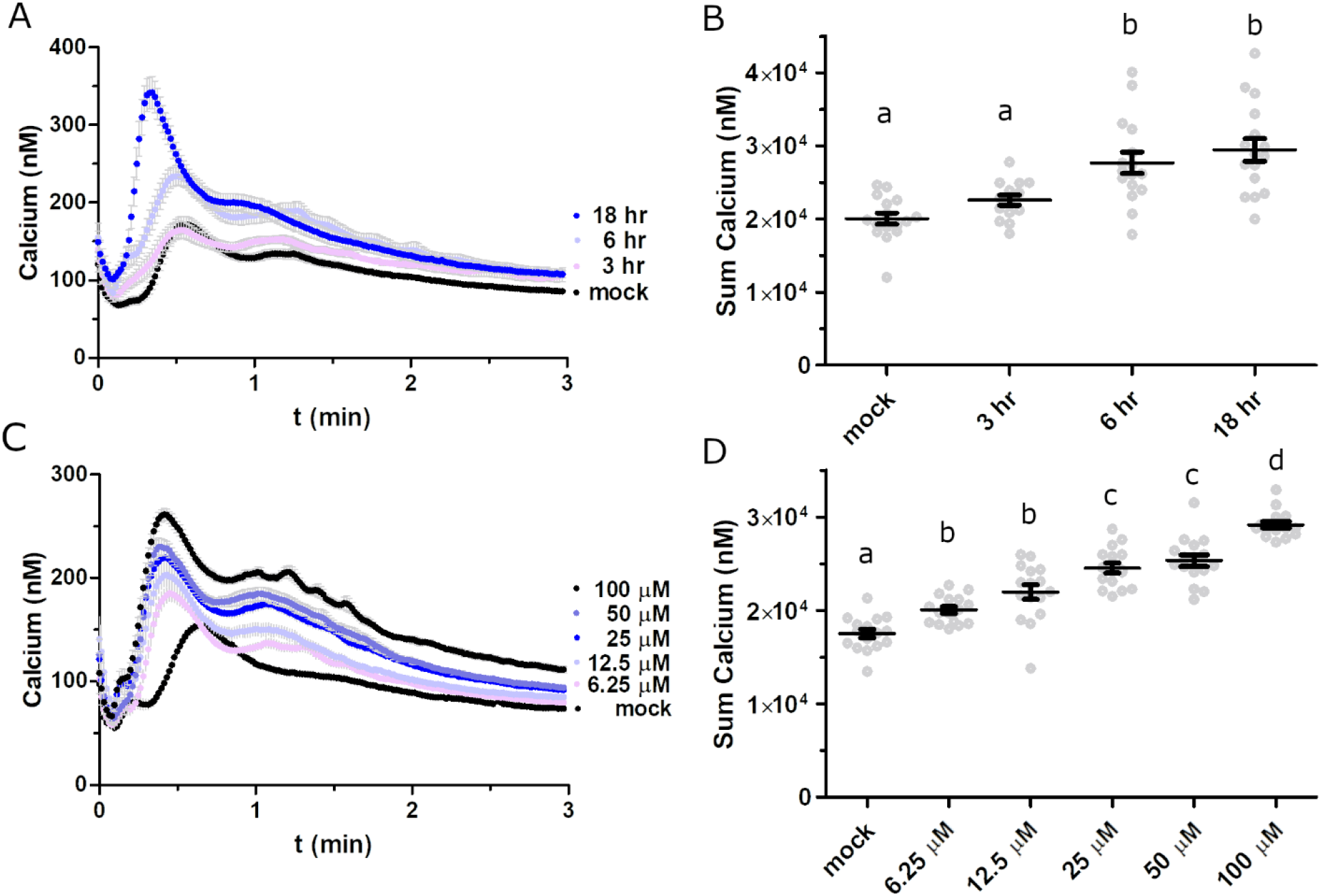
Effects of JA pre-treatment dosage and duration on ATP-induced calcium response. **(A)** Seedlings were treated as in Figure 1, with MeJA added to the reconstitution buffer 50 μM for 3, 6, or 18 hours, followed by recording of ATP-induced aequorin luminescence. **(B)** Summed calcium values from panel A. **(C)** Seedlings were treated as in Figure 1, with the reconstitution buffer containing varying MeJA concentrations. **(D)** Summed calcium values from panel C. All graphs show mean ± SEM (*n =* 14-16), with statistically significant differences indicated by different letters (*P* < 0.05, Tukey HSD).

### Differential effects of plant stress hormones on ATP-Induced cytosolic calcium response: enhancement by JA and suppression by SA

To examine whether other plant stress hormones impact eATP-induced [Ca^2+^]_cyt_ responses, we pre-treated AEQ seedlings overnight with either 50 μM MeJA, 300 μM salicylic acid (SA), 50 μM of the ethylene precursor aminocyclopropane-carboxylic acid (ACC), or 50 μM abscisic acid (ABA) prior to the ATP treatment. Consistent with previous results, MeJA pre-treatment significantly increased the ATP-induced [Ca^2+^]_cyt_ response by approximately 50% **(Figure 4)**. In contrast, the SA pre-treatment suppressed the ATP-induced [Ca^2+^]_cyt_ response by

**Figure 4.**
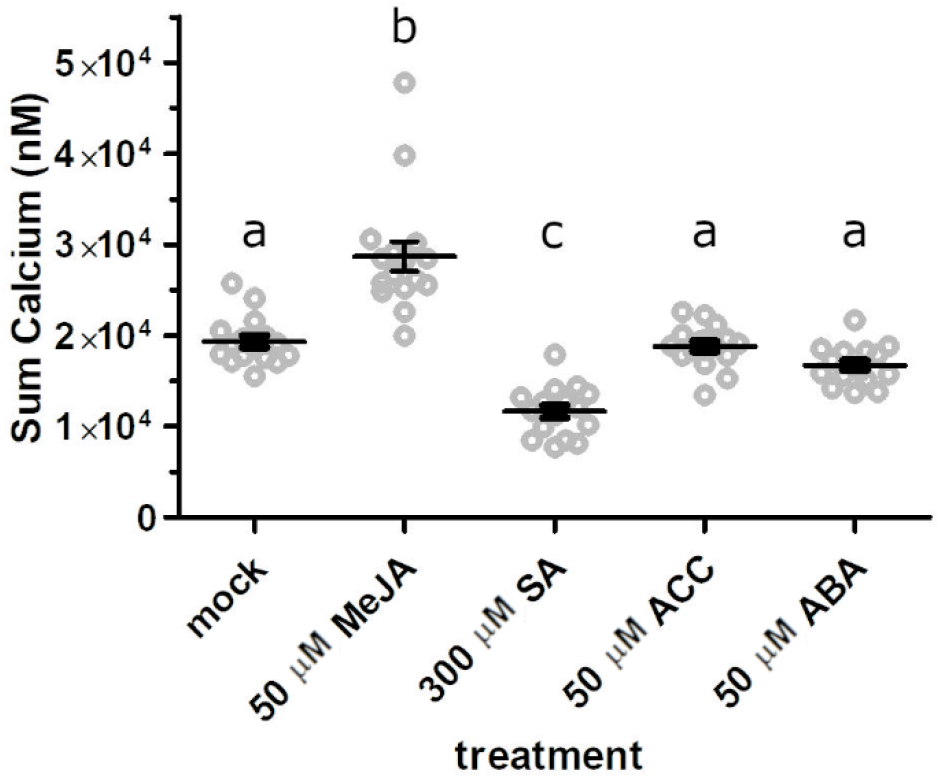
SA reduces ATP-induced calcium response, while JA increases it, but ABA and ACC have no effect. Aequorin-expressing seedlings were treated as in Figure 1, with overnight pre-treatment in reconstitution buffer containing 50 μM MeJA, 300 μM SA, 50 μM ACC, or 50 μM ABA, followed by recording ATP-induced luminescence. Data indicates summed calcium values over 2 minutes (mean ± SEM, *n =* 16 seedlings per treatment). Different letters indicate statistically significant differences (*P* < 0.05, Tukey HSD).

∼40% relative to mock-treated controls, while neither ABA nor ACC pre-treatment had a significant effect **(Figure 4)**. These findings suggest a unique role for JA in priming eATP signaling, while SA may act antagonistically modulating ATP-induced plant response in a distinct manner. These contrasting effects of MeJA and SA are consistent with extensive previous literature **(Van der Does et al., 2013)**.

### COI1 is essential for JA-primed eATP-induced cytosolic calcium response

Previous research on the interaction of JA and eATP signaling has primarily focused on transcriptional responses, without examining calcium signaling **(Jewell et al., 2019)**. To examine whether the ATP-induced [Ca^2+^]_cyt_ response relies on JA signaling, we crossed AEQ-expressing plants with *coi1-1* mutants to generate F3 lines segregating for the *coi1-1* mutant allele while maintaining AEQ expression. This segregating progeny approach is necessary given that homozygous *coi1* null mutants are male sterile **(Xie et al., 1998)**. AEQ seeds with *COI1-1*^*+/-*^ segregants were additionally exposed to 50 μM MeJA to differentiate *coi1-1* mutants from those with a WT *COI1* allele. After 7 days, these 3 genotypes were incubated overnight in coelenterazine reconstitution buffer with (all 3 genotypes) or without (WT) 50 μM MeJA and subsequently treated with 100 μM ATP. Consistent with our hypothesis, WT and *COI1*^+/-^ seedlings exhibited a robust ATP-induced [Ca^2+^]_cyt_ response following MeJA treatment **(Figure 5A, B)**. In contrast, the *coi1-1* mutant seedlings showed a significantly reduced ATP-induced [Ca^2+^]_cyt_ response, even after MeJA priming. To test the effects of COI1 mutation on basal and MeJA-primed eATP-responsive changes to [Ca^2+^]_cyt_ (i.e. without the MeJA pretreatment which is a necessity for selecting seedlings for treatment given the male sterility of COI1 null mutants), we used leaf discs from 4-week-old plants, *COI1*^*+/-*^ or *coi1-1*. Changes to [Ca^2+^]_cyt_ after ATP treatment were much lower in leaf discs relative to seedlings, and were not reliably observed in the absence of MeJA treatment **(Figure 5C,D)**. In addition, MeJA was unable to prime eATP calcium responses in the *coi1-1* mutant. These results indicate that COI1 is essential both for MeJA-induced priming of eATP-induced calcium response in seedlings and mature leaves, and for basal WT-like responses to eATP in seedlings.

**Figure 5.**
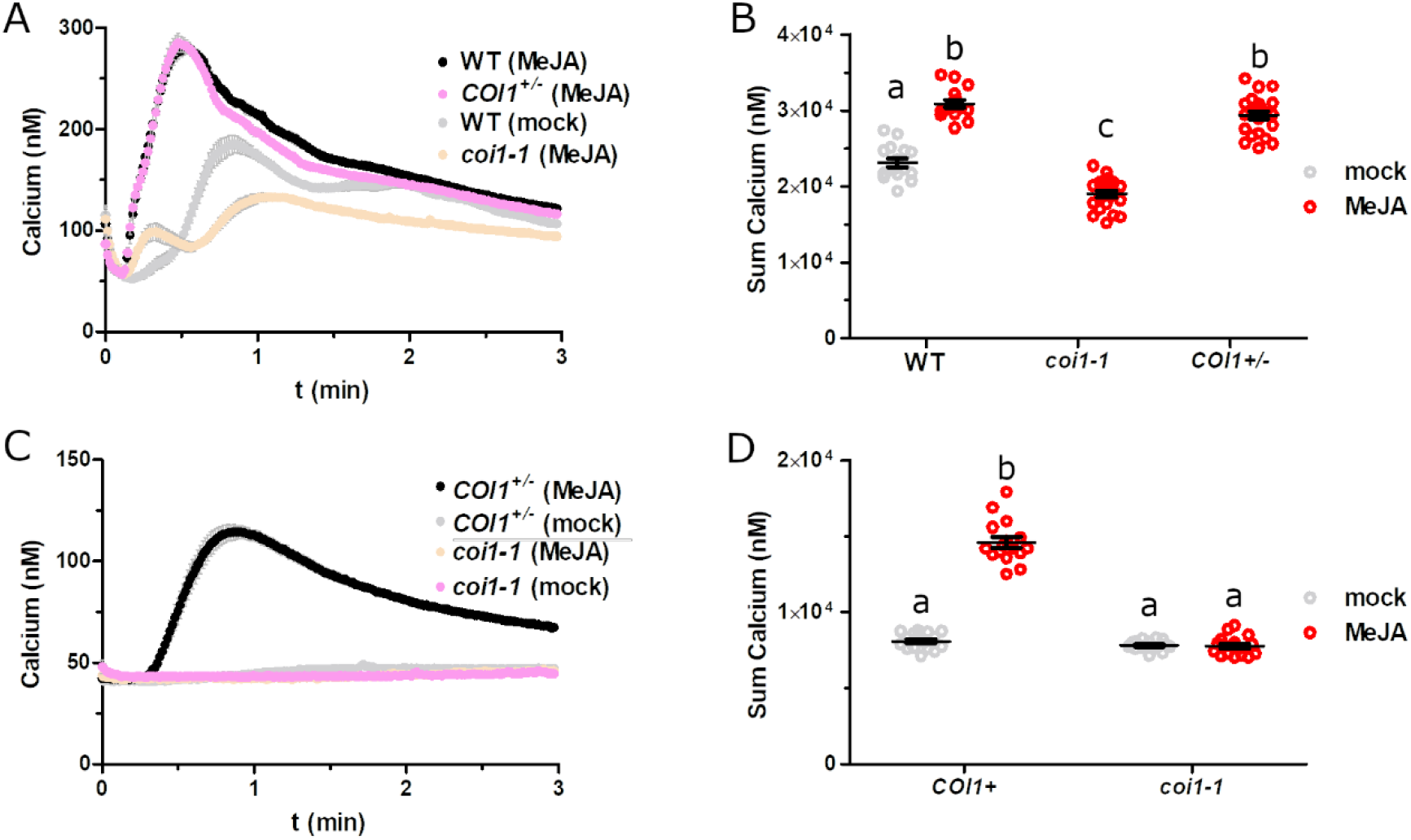
COI1 is required for both basal and MeJA-stimulated ATP-induced calcium response. (A) Seedlings of the indicated genotypes were treated overnight with 50 μM MeJA and 100 μM ATP-induced luminescence was recorded (mean ± SEM, *n =* 16-24). **(B)** Summed calcium values from panel A. Different letters indicate statistically significant differences (*P <* 0.05, Tukey HSD). **(C)** Leaf discs of the indicated genotype were treated overnight with or without 50 μM MeJA and 100 μM ATP-induced luminescence was recorded (mean ± SEM, *n =* 16). **(D)** Summed calcium values from panel C. Different letters indicate statistically significant differences (*P <* 0.05, Tukey HSD).

### *P2K1* gene expression is induced by JA treatment and wounding in a COI1-dependent manner

The eATP receptor P2K1 is known to be upregulated in response to wounding, with expression rising 2-to 3-fold in adult Arabidopsis within 1 hour and persisting for up to 24 hours **(Kilian et al., 2007)**. Similarly, treatment with 10 μM MeJA induces a ∼5-fold increase in *P2K1* transcript in seedlings within an hour **(Supplemental Figure S6A, Goda et al., 2008**; https://bar.utoronto.ca/efp/), and the bacterial JA-Ile mimic coronatine has been shown to elevate P2K1 expression in mature plants over the course of 24 hours **(Attaran et al., 2014)**. Additionally, chromatin immunoprecipitation sequencing experiments revealed that the JA-responsive transcription factors MYC2 and MYC3 bind to the presumptive *P2K1* promoter region **(Supplemental Figure S6B, Zander et al., 2020;** http://neomorph.salk.edu/MYC2). Furthermore, the *P2K1* transcript is induced significantly less (∼30% and 80% after 30 and 120 minutes, respectively) in the *myc2* mutant relative to WT after ∼5 μM gaseous MeJA treatment (**Supplemental Table S1**, Zander et al., 2020). Altogether, these previous data suggest that canonical JA signaling is likely required for JA-induced *P2K1* gene expression. To test this idea, we grew WT plants, *COI1*^*+/-*^ segregants, and the *coi1-1* mutants for four weeks, treated leaves 50 μM MeJA, and then quantified *P2K1* gene expression levels using RT-qPCR. As shown in **Figure 6**, MeJA treatment significantly increased *P2K1* expression by ∼6-fold in both WT and *COI1*^*+/-*^ plants, whereas no increase was observed in the *coi1-1* mutants. Similarly, 1 hr following wounding *P2K1* transcript levels significantly increased by ∼4-fold in WT and *COI1*^*+/-*^ plants but not *coi1-1* **(Figure 6B)**. This confirms that COI1 is required for *P2K1* induction in response to JA and wounding, correlating with the MeJA-potentiated and COI1-dependent increased [Ca2+]cyt in response to treatment with ATP.

**Figure 6.**
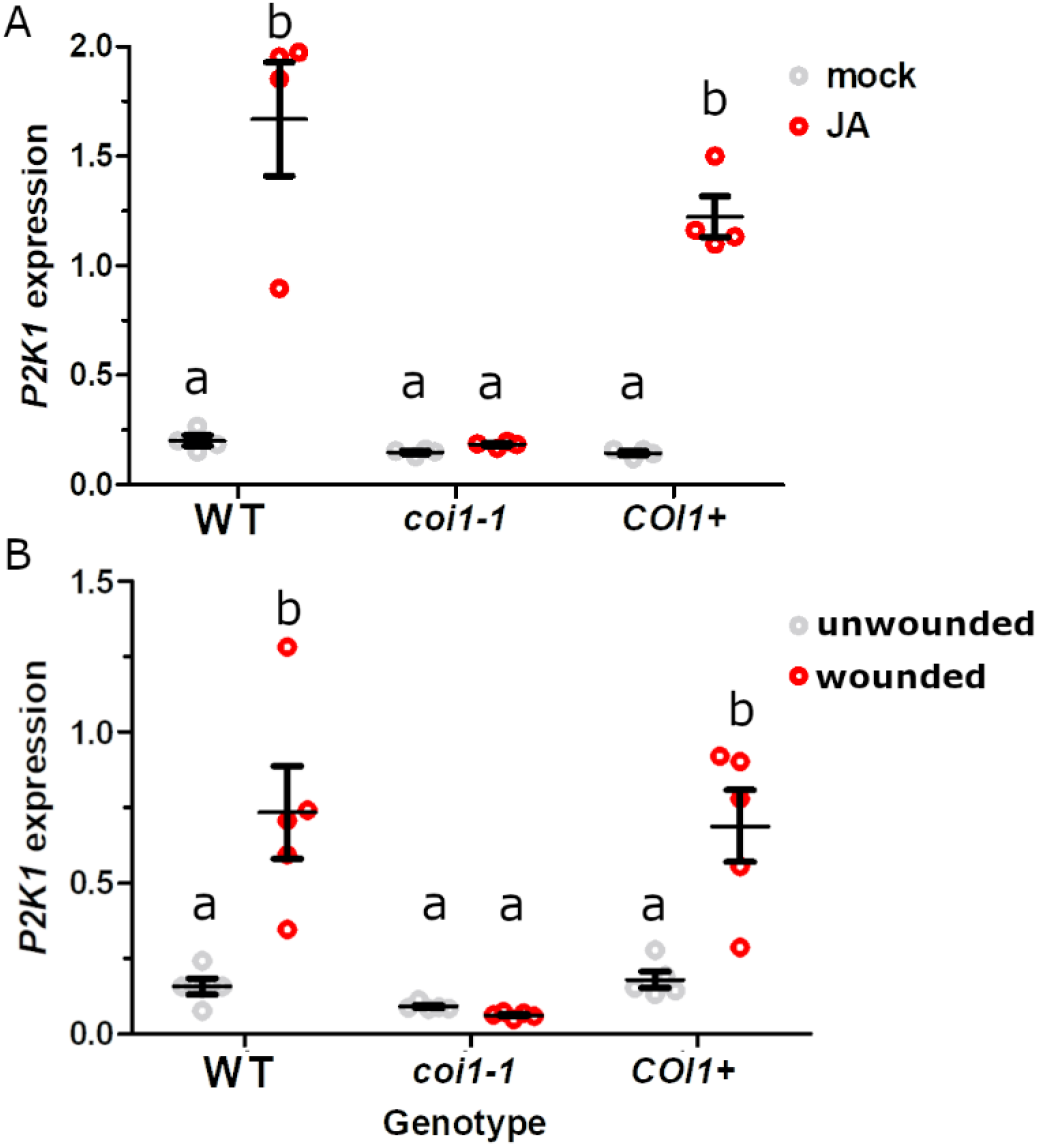
JA and wounding induces *P2K1* gene expression in a COI1-dependent manner. *P2K1* expression was measured in 4-week-old aequorin-expressing plants of the indicated genotypes either **(A)** 1 hour after a 50 μM MeJA spray or without MeJA treatment (n=4), or **(B)** 1 hour with or without wounding (n=5). Gene expression was normalized by *PP2A*. Mean, SEM, and individual values are shown. Different letters indicate statistically significant differences (*P* < 0.05, Tukey HSD).

### JA primes eATP-responsive gene expression

To further evaluate the physiological impact of JA priming on ATP responses, we tested whether MeJA pre-treatment enhances ATP-responsive gene expression. Given that many ATP-responsive genes also respond to JA **(Choi et al., 2014; Jewell et al., 2019)**, we treated plants with MeJA, then replaced the media to allow a 24-hour recovery period, thereby enabling JA-responsive genes to return to basal levels prior to the ATP treatment (see Materials and Methods). We selected eight genes with previously reported subtle ATP-induced expression changes, reasoning that an exaggeration of this marginal induction could be easier to detect. Across these genes, there were significant effects due to MeJA, ATP, and their interaction (*P*<2e-16, *P=*1.1e-10, and *P*=2.6e-3, respectively, two factor ANOVA F-test). Of these, six genes showed significant MeJA priming of ATP responsiveness (MeJA by ATP interaction, P<0.05, two factor ANOVA F-test): *LIPOXYGENASE 3* (*LOX3;* **Figure 7A**), *CALCIUM-DEPENDENT PROTEIN KINASE 28* (*CPK28*, **Figure 7B)**, *RESPIRATORY BURST OXIDASE HOMOLOGUE D* (*RBOHD*, **Figure 7C)**, *P2K1* (**Figure 7D)**, *JASMONATE-ZIM-DOMAIN PROTEIN 1* (*JAZ1*, **Supplemental Figure S7A)**, and *1-AMINOCYCLOPROPANE-1-CARBOXYLIC ACID SYNTHASE 6* (*ACS6*, **Supplemental Figure S7B)**. The remaining two genes, *RESPONSIVE TO DESSICATION 20* (*RD20*, **Supplemental Figure S7C)** and *MITOGEN-ACTIVATED PROTEIN KINASE 3* (*MPK3*, **Supplemental Figure S7D)**, did not show statistically significant MeJA priming. These findings suggest that JA priming enhances transcriptional responses to eATP, particularly for genes involved in stress and signaling pathways.

**Figure 7.**
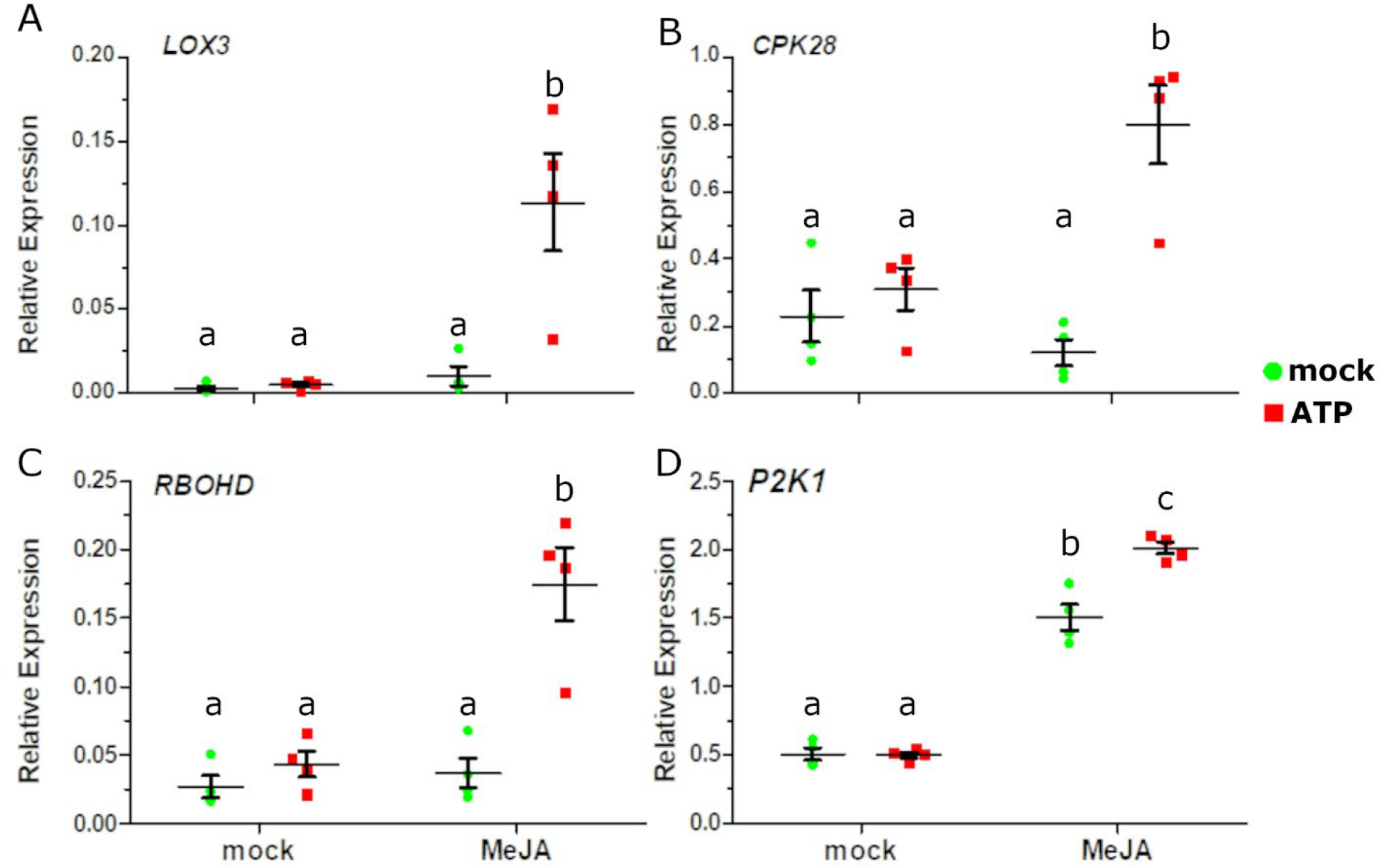
JA treatment potentiates eATP-induced transcriptional responses. Wild-type seedlings were pre-treated with or without 20 μM MeJA, followed by treatment with 0.5 mM ATP, as described in Materials and Methods. Gene expression was evaluated by RT-qPCR for **(A)** *LOX3*, **(B)** *CPK28*, **(C)** *RBOHD*, and **(D)** *P2K1*. Mean ± SEM and individual values are shown, with different letters indicating statistically significant differences in gene expression (*n* = 4, *P* < 0.05, Tukey HSD).

## DISCUSSION

In this study, we report that JA potentiates two Arabidopsis responses to eATP through mechanisms dependent on the JA receptor COI1 and the eATP receptor P2K1. Our findings further indicate that JA treatment induces *P2K1* gene expression in a COI1-dependent manner, suggesting that JA-primed eATP response acts by transcriptional upregulation of *P2K1*. Supporting this, published ChIP-seq data **(Zander et al., 2020)** demonstrate specific binding of MYC2 and MYC3 transcription factors to the *P2K1* promoter, suggesting that JA induction of *P2K1* relies on the canonical JA signaling pathway involving bHLH MYC transcription factors. Importantly, the priming effect was observed even at low JA concentrations, consistent with endogenous JA levels after wounding **(Kimberlin et al., 2022)**, hinting at the physiological relevance of this JA-eATP signaling interaction. Interestingly, we also observed that MeJA treatment of Arabidopsis leaf discs induced ATP release into surrounding media. Though the physiological relevance and mechanistic basis of this observation is unknown, it suggests multiple levels to the JA stimulation of eATP signaling.

Since the initial report of ATP-induced degradation of JAZ1 in 2018 **(Tripathi et al.)**, no studies have corroborated or explained this phenomenon. While JAZ1 degradation is not central to our conclusions, its irreproducibility has implications for understanding the full mechanism by which eATP activates JA-responsive genes. It remains possible that other JAZ proteins could play a role in eATP-induced activation of JA-responsive genes. For example, JAZ8, which lacks a canonical degradation mechanism within JA signaling (**Shyu et al., 2012**) regulates both JA-responsive and JA biosynthesis genes in response to stress-induced [Ca^2+^]_cyt_ elevations (**Yan et al., 2018)**, a hallmark of eATP response. Despite the uncertainty surrounding JAZs, activation of JA-responsive genes in this pathway clearly relies on the P2K1 receptor and the MYC and CAMTA3 transcription factors **(Jewell et al., 2019)**, with upstream signaling involving the P2K1-interacting and -phosphorylated INTEGRIN-LINKED KINASE 5 (ILK5, **Kim et al, 2023b**). Recently, it was suggested that phosphorylation of ZmMYC2 transcription factors by the maize mitogen-activated protein kinase (MAPK) ZmMPK4 enhances transcriptional activation activity of ZmMYC2s **(Li et al, 2024)**. Given that MPK phosphorylation is a characteristic feature of eATP responses **(Choi et al, 2014; Cho et al, 2022)** and is reduced in *ilk5* Arabidopsis mutants **(Kim et al, 2023b)**, these cross-species comparisons suggest that eATP may activate MYC2 through a MAPK cascade. Experimental evaluation of this model should be an interesting direction for future research. Collectively, these findings suggest that JA and eATP pathways amplify each other’s signaling as part of a defense priming mechanisms during stress responses.

Given the roles of JA and eATP signaling in responses to biotic stresses such as pathogens and herbivores **(Browse 2009; Chivasa et al., 2009)**, it is attractive to speculate that this interaction promotes heightened defense responses after initial injury. However, a novel role for JA signaling in wound healing and tissue regeneration has recently emerged **(Zhang et al., 2019; Zhou et al., 2019)**. In addition, the wound-induced tomato elicitor peptide REF1, an orthologue of Arabidopsis PLANT ELICITOR PEPTIDEs (PEPs), dramatically increased wound-induced callus formation and shoot regeneration **(Yang et al., 2024)**, and exogenous ATP treatment induces expression of several PEP-encoding genes (*PROPEP1, PROPEP3, PROPEP4*, and *PROPEP7*) and two PEP receptor genes in a P2K1-dependent manner **(Jewell et al., 2019)**. While Arabidopsis has eight PEPs, to our knowledge none have been investigated for a tissue regeneration activity similar to REF1. This raises the question: does JA potentiation of eATP response contribute not only to defense priming but also to tissue regeneration after wounding? This hypothesis opens exciting avenues for further research. Given the well-documented role of JA in pathogen defense and the more nuanced roles associated with eATP signaling, investigating the potential role of JA-eATP interaction in tissue regeneration may provide new insights into wound response mechanisms.

## MATERIALS AND METHODS

### Plant Materials

Arabidopsis (*Arabidopsis thaliana)* WT AEQ (pMAQ2.4), *p2k1-3 AEQ, 35S-JAZ1-GUS, aos 35S-JAZ1-GUS* and *coi1-1* were described previously **(Knight et al., 1991; Choi et al., 2014; Thines et al., 2007; Tripathi et al., 2018; Xie et al., 1998)**. Columbia-0 (CS70000) was obtained from the Arabidopsis Biological Resource Center (https://abrc.osu.edu/). Due to the male sterility of *coi1* null alleles, *COI1-1*^*+-*^ *AEQ* lines were maintained as segregating for the *coi1-1* mutation but fixed for AEQ expression. These were identified by crossing WT AEQ and *coi1-1*, selfing F2 plants and selecting F3 lines based on segregating MeJA insensitivity andfixed 25 μg/mL kanamycin resistance. AEQ expression was verified based on luminescence in the AEQ discharge solution **(Choi et al., 2014)**.

### Reagents

All chemicals except as otherwise indicated were obtained from Sigma-Aldrich. ATP was prepared as a 0.1 M stock solution in 50 mM MES buffer (pH 5.7) and adjusted to pH 5.7 using sodium hydroxide. Methyl jasmonate (Bedoukian Research), salicylic acid, and abscisic acid were prepared in DMSO as 0.1 M, 0.3 M and 10 mM stock solutions, respectively. 1-aminocyclopropane-1-carboxylic acid was dissolved in distilled deionized water at 0.1 M. Coelenterazine native (NanoLight Technology) was prepared as a 1 mM stock solution in 100% ethanol (v/v). All these solutions were stored at –20°C in a sealed, dark container to prevent degradation.

### Assays to Monitor Changes in Cytosolic Calcium

Assays to monitor changes in [Ca^2+^]_cyt_ were performed as described previously **(Choi et al., 2014)**. Briefly, surface-sterilized and cold-stratified seeds were sewn to square plates containing ½-strength Murashige and Skoog (MS) media (pH 5.8) with or without 50 μM MeJA and with 0.8% (w/v) sucrose, 0.8% (w/v) agar, and 0.5g/L MES pH-adjusted using potassium hydroxide. After sowing, plates were transferred to a 22°C growth chamber (Conviron) with 12-h light cycle (100-120 μmol photons m^-2^ s^-1^) and grown vertically for 7 days. On the morning of the 8^th^ day, individual seedlings were carefully transferred to single wells of white 96-well plates containing 50 μL aequorin reconstitution buffer containing 10 μM coelenterazine (NanoLight Technology) with or without various hormones as described in the Results section, 2 mM MES buffer pH 5.7, and 10 mM CaCl_2_-heptahydrate and incubated in darkness overnight. For 50 μM MeJA treatments, the final concentration of DMSO in the treatment solution was 0.05%. Luminescence was monitored after adding 50 μL of 2X treatment solution via autoinjector using the GloMax Navigator (Promega). After elicitor-induced luminescence was recorded, 100 μL of discharge solution (20% ethanol (v/v) and 2M CaCl_2_-heptahydrate) was added to each well, and luminescence recorded for an additional 30 s. Luminescence data were converted to [Ca^2+^]_cyt_ using the formula of **Knight et al., 1996**. Summed calcium data in figures were calculated by adding [Ca^2+^]_cyt_ values at each timepoint.

To test cytosolic calcium responses to eATP of *COI1*^*+/-*^ versus *coi1-1* absent MeJA pretreatment, seedlings were grown and selected as above and then transferred to soil in a 22°C growth chamber (Conviron) with 12-h light cycle (100-120 μmol photons m^-2^ s^-1^). After ∼4 weeks, leaf discs from expanded leaves were collected with a 4 mm biopsy punch (Ted Pella, Inc.) into distilled water in white 96-well plates and allowed to sit under light for 1 hour. Thereafter, the water was removed and 50 μL aequorin reconstitution buffer was added containing either 50 μM MeJA or 0.05% DMSO and plates moved to darkness overnight (∼18 hours) after which luminescence was recorded as above.

### Quantitative JAZ1-GUS Stability Assay

The quantitative GUS assay was performed as described **(Jefferson et al., 1987)**. Five to ten 10-day old seedlings of *35S-JAZ1-GUS* **(Thines et al., 2007)** or *aos 35S-JAZ1-GUS* were incubated overnight in buffer (10 mM CaCl_2_, 2 mM MES, pH5.7). The next day, seedlings were treated with buffer alone (mock), 1 mM ATP, or 2 μM MeJA for 30 minutes in the light, then blotted dry and snap-frozen in liquid nitrogen. These concentrations are on the order of concentrations described in the original manuscript (Tripathi et al., 2018). The seedlings were ground in 75 μL of GUS extraction buffer (GEB; 50 mM sodium phosphate pH 7.0, 10 mM EDTA, 10 mM β-mercaptoethanol, 0.1% triton X-100, 0.1% sarkosyl) and centrifuged at 4°C for 15 minutes at 15,000 rcf to remove insoluble debris. Supernatant (65 μL) was collected to separate tubes on ice and protein quantity estimated using Bradford reagent. For each sample, equal amounts of protein (∼9 μg) were added to individual wells of a white 96-well plate in a final volume of 100 μL GEB containing 1 mM 4-Methylumbelliferyl-β-D-glucuronide hydrate (MUG) prewarmed to 37°C after which the plate was stored in a dark 37°C incubator. At 60 minutes, 60 μL aliquots were added to 60 μL of 0.2 M Na_2_CO_3_ stop buffer in a separate white 96-well plate. Fluorescence from MU released in each well was measured in a plate reader with excitation at 365 nm and emission at 455 nm, and rate was calculated as light units/mg/min. In preliminary experiments, signal was not reliably detected at 15 minutes incubation, but activity was linear between 30 to 90 minutes.

### Quantification of Extracellular ATP

Extracellular ATP was quantified as previously described **(Ramachandran et al., 2019)** using Col-0 plants one month after sewing. Individual leaf discs (4 mm, Electron Microscopy Sciences) were collected into 150 μL of distilled deionized water in individual wells of a 96 well plate and the plate transferred to a growth chamber. Two hours later, 50 uL of water, 0.8 M NaCl, 0.2% DMSO, or 200 μM MeJA were added to each well. At 15, 60, and 960 minutes after addition of these solutions, 50 μL was gently removed from each well and immediately flash frozen. Thereafter, these were heated to 96° C for 5 minutes. After allowing to cool, 25 μL was added to wells of a white 96 well plate, and 25 μL of luciferase/luciferin reagent (ENLITEN, Promega) was added to each well by the luminometer’s autoinjector and luminescence recorded for 5 seconds. A standard curve was fit by use of 100, 50, 33, 10, 1, 0.1, and 0 nM ATP to calculate ATP concentrations in the wells.

### RT-qPCR Analysis

For MeJA-primed eATP-induced gene expression experiments, Col-0 seedlings were grown as described above. On the 7^th^ day, 20 seedlings were transferred to 6-well plates containing 2 mL of ½-strength sterile MS liquid media (without sucrose and agar) with or without 20 μM MeJA and returned to the growth chamber. The next day, the MeJA-containing liquid media was removed, and 2 mL of fresh liquid media was added to each well. Finally, on the 9^th^ day, 2 mL of liquid media with or without 0.5 mM ATP was added, a concentration optimized previously (Jewell et al., 2019), and the plates were returned to the growth chamber for 30 minutes. Thereafter, seedlings were removed from the wells, gently blotted on tissue paper, flash frozen in liquid nitrogen, and stored for the subsequent RT-qPCR experiments. All media changes were performed in a sterile laminar flow hood.

For the MeJA-induced *P2K1* gene expression studies, WT AEQ, and AEQ lines segregating for the *coi1-1* mutation were initially grown as above for 7 days, then 8 (WT) or ∼60 (*COI1*^*+/-*^) seedlings were transferred to autoclave-sterilized soil in a growth room (22°C, 12 h light, 100-120 μmol photons m^-2^ s^-1^) and grown for an additional 3 weeks. Adult plants were used so that we could reliably determine the genotype of plants without prior MeJA treatment. A single leaf from each plant was excised with a scalpel and snap frozen and stored. Immediately after initial harvest, all plants were uniformly sprayed with 50 μM MeJA in 0.1% Tween-20 until they were dripping. After 1 hour, another set of single leaves were harvested, snap-frozen, and stored at -80°C. Thereafter, the same plants were sprayed twice over 2 weeks with 500 μM MeJA in 0.1% Tween-20, which induced anthocyanin accumulation in plants with a functional *COI1* allele. After flowering, sterile plants (a hallmark of *coi1* null mutants) were noted. Sterile plants not accumulating anthocyanin were recorded as *coi1-1* mutants. Leaves of four plants from each genotype (AEQ, *coi1-1*, and *COI*^*+/-*^) were selected for subsequent RT-qPCR experiments. Plants for wounding and RT-qPCR were grown similarly. Individual leaves of individual plants were wounded twice across the midvein with a hemostat, damaging ∼40-50% of leaf area. One hour later, 2 leaves from separate plants (wounded or unwounded) were combined as a replicate with a total of 5 replicates for each genotype/treatment combination.Frozen plant tissue was disrupted with a bead beater (MIni-Beadbeater-96, BioSpec Products) and total RNA was isolated using the Quick-RNA Miniprep Kit (Zymo Research), including the on-column DNase I treatment as per the manufacturer’s instructions, which we have found is critical for reproducible RT-qPCR results with low-expressed eATP responsive genes.

After RNA isolation, 1 μg of RNA was used in a 20-μL reverse transcription reaction according to the manufacturer’s instructions (iScript; Bio-Rad) followed by a 5-fold dilution with ultrapure water. RT-qPCR was performed using 2 μL of diluted cDNA in a 20-μL reaction with SYBR Green dye polymerase mix (SsoAdvanced; Bio-Rad) in white-walled plates in a CFX96 thermocycler (Bio-Rad). Gene expression was normalized to the reference gene *PP2A* (AT1G13320) as described **(Czechowski et al., 2005; Rieu and Powers 2009)**. Primers used are listed in **Supplemental Table S2**.

### Statistical Analyses

Figures were created using GraphPad Prism 5 (GraphPad Software). Statistical analyses were performed in R version 4.3.1 **(R CoreTeam, 2020)**, using ANOVA with Tukey’s HSD post-hoc testing where appropriate.

### Accession Numbers

Sequence data and gene information from this article can be found at The Arabidopsis Information Resource (https://www.arabidopsis.org/).

## DATA STATEMENT

Seeds and data underlying the figures are available upon request from Jeremy Jewell or Kiwamu Tanaka.

## ACKNOWLEDGEMENTS

Thank you to John Browse for the *35S-JAZ1-GUS* seeds, the *coi1-1* seeds, the *aos* seeds, leadership by example, and support and encouragement throughout the years. Thank you to Marc Knight for the pMAQ2.4 seeds. Thank you to Nathan Havko for insightful discussion. This work was funded by the NSF (no. IOS-1557813, IOS-2048410) and USDA NIFA (Hatch project no. 1015621). Funding was also provided by the Center for Bioenergy Innovation (CBI) led by Oak Ridge National Laboratory (ORNL). CBI is funded as a DOE Bioenergy Research Center supported by the BER Program in the DOE Office of Science under FWP ERKP886. ORNL is managed by UT-Battelle, LLC, for the DOE under contract no. DE-AC05-00OR22725.

## AUTHOR CONTRIBUTIONS

JBJ designed experiments with input from KT. JBJ, AC, and JPT performed experiments. JBJ performed statistical analyses of data. JBJ, KT,and AC wrote the paper with suggestions and approval of all authors.

## SHORT LEGENDS FOR SUPPORTING INFORMATION

**Supplemental Figure S1. Exogenous ATP treatment does not reduce JAZ1 protein stability in *aos* mutant seedlings, while MeJA does**.

**Supplemental Figure S2. The time to maximal ATP-induced [Ca**^**2+**^]_**cyt**_ **response is reduced after MeJA treatment**.

**Supplemental Figure S3. Effect of overnight DMSO treatment on ATP-induced changes in [Ca**^**2+**^]_**cyt**_.

**Supplemental Figure S4. Effect of water, DMSO, or MeJA on [Ca**^**2+**^]_**cyt**_.

**Supplemental Figure S5. Effect of NaCl or MeJA on ATP release from leaf discs**.

**Supplemental Figure S6 Induction of *P2K1* by JA may be mediated by MYC transcription factors**.

**Supplemental Figure S7. JA-primed eATP-responsive gene expression**.

**Supplemental Table S1. Reduced induction of *P2K1* gene by MeJA treatment in the *myc2* mutant**.

**Supplemental Table S2. Primers used in this study**.

