## Supporting Information for "Jasmonate Primes Plant Responses to Extracellular ATP"

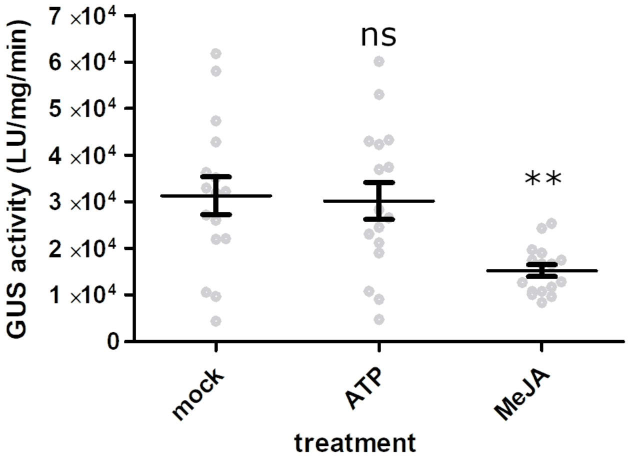


**Supplemental Figure S1. Exogenous ATP treatment does not reduce JAZ1 protein stability in *aos* mutant seedlings, while MeJA does.** Mutant *aos* seedlings expressing JAZ1-GUS were treated with 2 μM MeJA or 1 mM ATP for 30 minutes and processed for quantitative GUS activity measurements to assess JAZ1 protein stability as described in Methods. Statistical comparisons to mock-treated seedlings are indicated by ** (*P* < 0.01) and ns (*P* = 0.97, Fisher's LSD). Error bars indicate SEM, *n*=16 replicates per treatment across 2 trials.

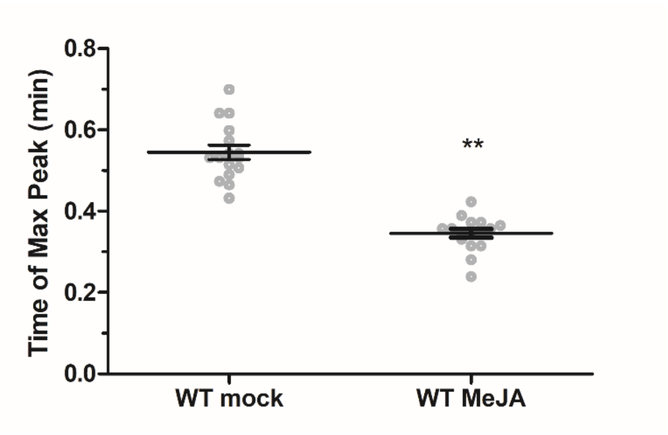


**Supplemental Figure S2. The time to maximal ATP-induced $[Ca^{2+}]_{cyt}$ response is reduced after MeJA treatment.** Calcium values from Figure 2B were manually inspected for the time of maximum calcium level. MeJA pretreatment resulted in ~37% earlier maximal calcium response in WT seedlings (*** *P* < 0.001, two-sided t-test). Error bars indicate SEM, *n*=16.

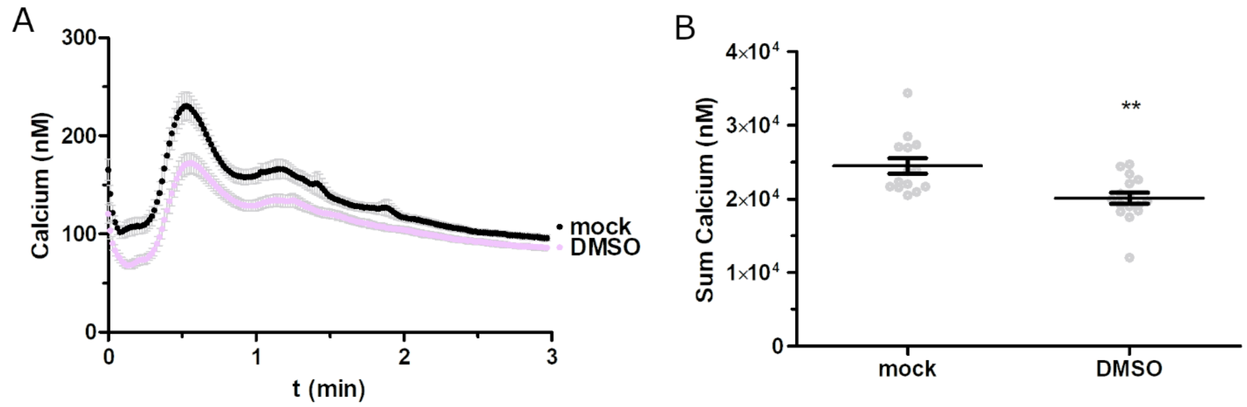


**Supplemental Figure S3. Effect of overnight DMSO treatment on ATP-induced changes in $[Ca^{2+}]_{cyt}$. (A)** Seedlings were incubated in reconstitution buffer with or without 0.05% DMSO overnight and treated with 100 µM ATP and aequorin luminescence was recorded for 3 minutes. **(B)** Summed calcium values from panel A. DMSO treatment resulted in ~18% reduction in calculated calcium values (** $P < 0.01$, two-sided t-test). Error bars indicate SEM, $n$=14-16 seedlings.

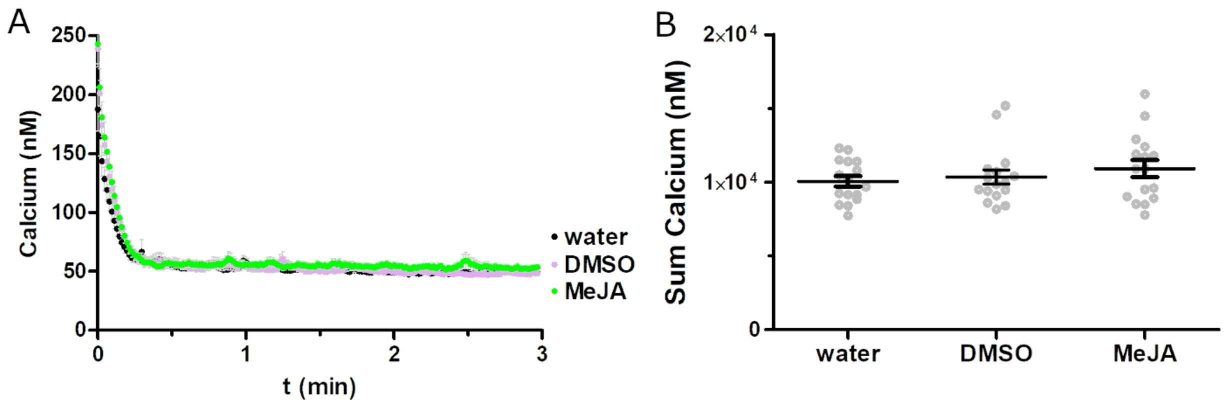


**Supplemental Figure S4. Effect of water, DMSO, or MeJA on $[Ca^{2+}]_{cyt}$. (A)** Seedlings were incubated overnight in reconstitution buffer then treated with water, 0.05% DMSO, or 50 µM MeJA and aequorin luminescence was recorded for 3 minutes. **(B)** Summed values from panel A. There were no significant differences in total calcium levels between the treatments ($P > 0.4$, one factor ANOVA). Error bars indicate SEM, $n$=16 seedlings.

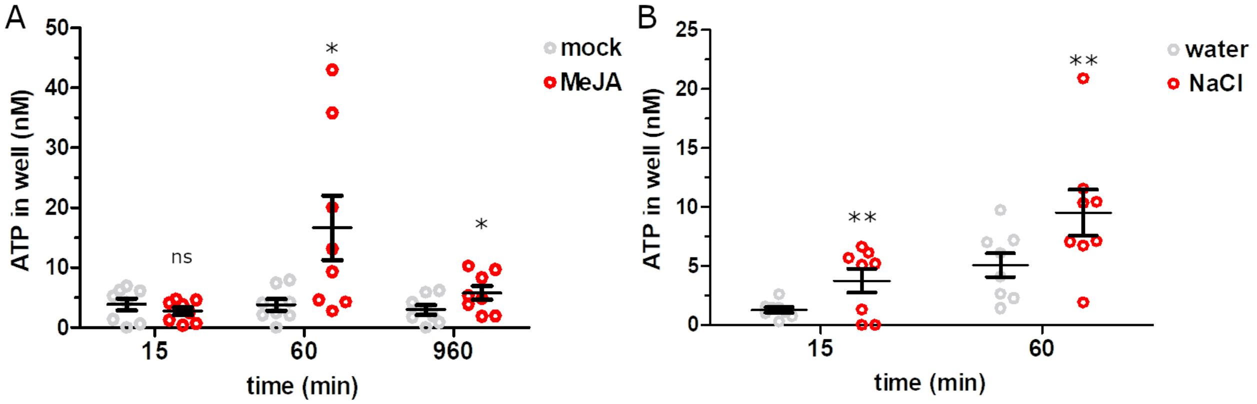


**Supplemental Figure S5. Effect of NaCl or MeJA on ATP release from leaf discs.** Leaf discs were treated at time 0 with **(A)** water or 200 mM NaCl, or **(B)** 0.05% DMSO or 50 μM MeJA and incubation media was collected at the indicated time points and assayed for ATP concentration. Error bars indicate SEM, *n*=8 leaf discs. (ns $P > 0.1$, * $P < 0.05$, ** $P < 0.01$, 1-sided t-test.)

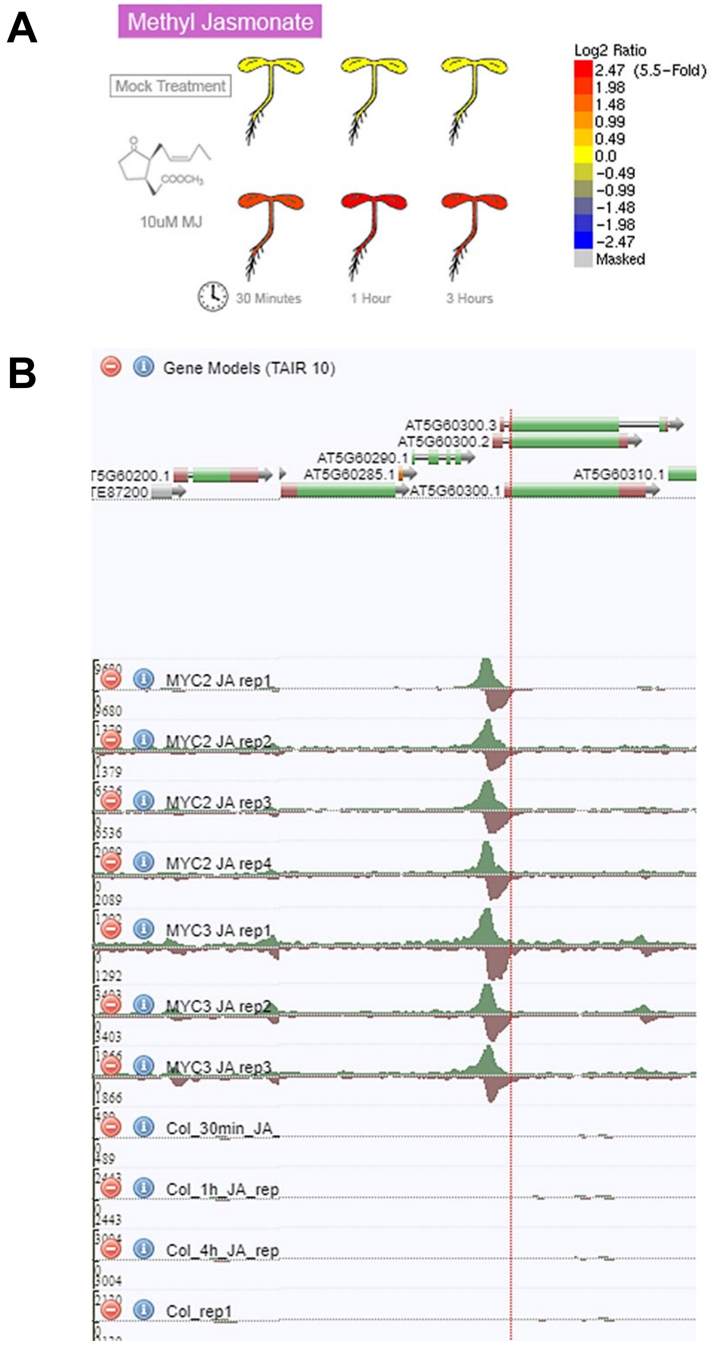


**Supplemental Figure S6. Induction of P2K1 by JA may be mediated by MYC transcription factors. (A)** Data from the Arabidopsis eFP Browser (Winter et al., 2007) shows strong *P2K1* upregulation by MeJA treatment, visualized at http://bar.utoronto.ca/efp/cgi-bin/efpWeb.cgi. **(B)** ChIP-seq data (Zander et al., 2020) demonstrate specific MYC2 and MYC3 binding to the promoter region of *P2K1* (AT5G60300), visualized at http://neomorph.salk.edu/MYC2. Note that the Y-axis for MYC data is on the order of thousands of reads, while the Col-0 data is on the order of 100s.

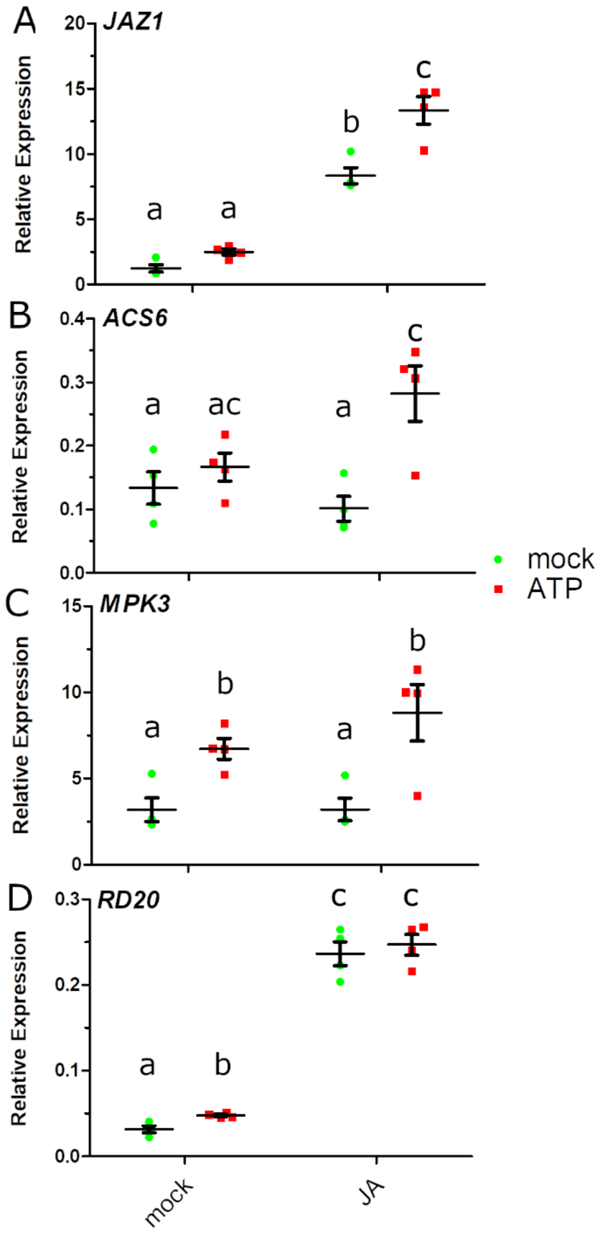


**Supplemental Figure S7. JA-primed eATP-responsive gene expression.** Wild-type seedlings were pre-treated with or without 20 µM MeJA, followed by 0.5 mM ATP treatment as described in Materials and Methods. Gene expression was evaluated by RT-qPCR for **(A)** *JAZ1*, **(B)** *ACS6,* **(C)** *RD20*, and **(D)** *MPK3*. Mean ± SEM and individual values are shown with different letters indicating statistically significant differences in gene expression ($n$ = 4, $P < 0.05$, Tukey HSD).

**Supplemental Table S1. Reduced induction of *P2K1* gene by MeJA treatment in the *myc2* mutant.** Data from Zander et al., 2020.

| Time after JA treatment (minutes) | Fold change of *P2K1* expression (Col-0/*myc2*) | FDR |
|---|---|---|
| 0 | 1.15 | 0.46 |
| 30 | 1.30 | 0.04 |
| 120 | 1.78 | 1.2e-5 |
| 240 | 0.96 | 0.86 |

**Supplemental Table S2. Primers used in this study.**

| Primer | Sequence 5'-3' | Reference |
|---|---|---|
| qJAZ1 f | GAGCAAAGGCACCGCTAATA | Grunewald et al. 2009 |
| qJAZ1 r | TGCGATAGTAGCGATGTTGC | |
| qMPK3 f | GCTTGGCACACCGACAGAATC | Choi et al. 2014 |
| qMPK3 r | CGTGGGAAGTTGGGAAGTTGC | |
| qP2K1 f | TGGAGTTTGTCAGGTCCATCG | Choi et al. 2014 |
| qP2K1 r | CTGAGGATCTTCTGCAGGCAA | |
| qPP2A f | TAACGTGGCCAAAATGATGC | Czechowski et al. 2005 |
| qPP2A r | GTTCTCCACAACCGCTTGGT | |
| qRD20 f | TGACACCGAAGGAAGGTATGTCC | Choi et al. 2014 |
| qRD20 r | CTTTAACCGTTAGCGCGTATTTGC | |
| qCPK28 f | ACCCACGAGCACGGCTAA | Choi et al. 2014 |
| qCPK28 r | TTCTCTAACCCACGCATGTGAT | |
| qRBOHD f | CATGCGGGTGCCCATTT | Choi et al. 2014 |
| qRBOHD r | ATCCGCGGCAATTAAACG | |
| qLOX3 f | GCTCGCTAAAGCCCACGTTAGTTC | Choi et al. 2014 |
| qLOX3 r | AGCATGCATGTGTCCGTAACCAG | |
| qACS6 f | CAGCAACGTTTGATTCGGAAA | Choi et al. 2014 |
| qACS6 r | CGTTGAGCTTCACTTGGTGAAC | |

Winter D, Vinegar B, Nahal H, Ammar R, Wilson GV, Provart NJ. An "electronic fluorescent pictograph" browser for exploring and analyzing large-scale biological data sets. PloS ONE 2(8):e718
